# MisMatch Negativity-study showing pre-lexical sensitivity to both primary Final Accent and secondary Initial Accent in French

**DOI:** 10.1101/727776

**Authors:** Noémie te Rietmolen, Radouane El Yagoubi, Corine Astésano

## Abstract

In French, accentuation is not lexically distinctive and tightly intertwined with intonation. This has led to the language being described of as ‘a language without accent’ and to French listeners being alleged ‘deaf to stress’. However, if one considers Di Cristo’s model in which the metrical structure of speech plays a central role, it becomes possible to envision stress templates underlying the cognitive representation of words. This event-related potential (erp) study examined whether French listeners are sensitive to the French primary final accent (fa) and secondary initial accent (ia), and whether the accents are part of the French phonologically expected stress pattern. Two oddball studies were carried out. In the first study, in one condition, deviants were presented without (−fa) and standards with final accent (+fa), while in another condition, these positions were switched. We obtained asymmetric mmn waveforms, such that deviants −fa elicited a larger mmn than deviants +fa (which did not elicit an mmn), pointing toward a preference for stress patterns with fa. Additionally, the difference waveforms between identical stimuli in different positions within the oddball paradigms indicated −fa stimuli to be disfavored whether they were the deviants or the standards. In the second study, standards were always presented with both the initial and final accent, while deviants were presented either without final accent (−fa) or without initial accent (−ia). Here, we obtained mmns both to deviants −fa and to deviants −ia, although −fa deviants elicited a more ample mmn. Nevertheless, the results show that French listeners are not deaf to the initial and final accents, pointing instead to an abstract phonological representation for both accents. In sum, the results argue against the notion of stress deafness for French and instead suggest accentuation to play a more important role in French speech comprehension than is currently acknowledged.

## 1 Introduction

French accentuation holds a low phonological and post-lexical status, i.e. it is not considered to directly apply to the word domain, but, instead, held to belong to the phrase. This means that the accents have post-lexical functions, and are not thought to contribute to word processing. For instance, French accents may signal phrase boundaries or present the utterance’s information structure, but they can never distinguish the semantic content of a word. Two (surface-level) group accents are generally recognized in French, the final accent (fa) and the initial accent (ia). fa is the primary stress, obligatory marking the right boundary of the accentual phrase (ap; Jun & Fougeron, 2000) with a lengthened syllable rime, sometimes supported by an additional fluctuation in *f*_0_. This accent is the compulsory accent in French and falls on the last syllable of the last word of ap, i.e. fa typically co-occurs with the right prosodic constituent boundary. The second accent, ia, is the secondary stress, optionally marking the left boundary of ap. This accent is primarily cued by a rise in *f*_0_ and a secondary lengthening of the syllabic onset (Astésano, 2001). ia is mostly associated with its rhythmic function, i.e. it intervenes when a long stretch of syllables is pronounced without fa (a so-called stress lapse). French accentuation is thus not lexically distinctive and tightly intertwined with post-lexical, intonational prominence. These two properties of accentuation have led the existence of the accent being questioned for French (Rossi, 1980), and to the notion of French as *a language without accent* becoming the generally accepted view on French prosody, such that accentuation is attributed a rather trivial role in speech processing. Indeed, as some authors have argued, if French language does not know lexical stress, it is reasonable to assume that its speakers are confronted with stressed syllables too infrequently to be able to hear the accents (e.g. Dupoux et al., 1997). That is, the rare interactions with local prominences in ‘a language without accent’ are presumed insufficient for speakers to develop a sensitivity to accentual information, essentially leaving them ‘deaf to stress’.^1^ Because listeners can still readily decode speech, despite their supposed ‘phonological deafness’, it—according to these scholars—stood to reason that accentuation is unlikely to play an important function in French comprehension processes. Consequently, French accentuation has attracted rather little interest in the linguistic field.

However, if one considers Di Cristo’s model in which the metrical structure of speech plays a central role (Di Cristo, 2000), it becomes possible to envision the accents to be encoded in stress templates underlying the cognitive representation of the lexical word. In turn, if stress templates are phonologically encoded at the level of the word, they may readily contribute to speech comprehension. Studies investigating the phonological status of French accentuation have indeed reported results in favor of a sensitivity to the metrical structure of words. Not only were metrical incongruences (stress on the medial syllable, a violation in French) found to slow down semantic processing (Magne et al., 2007), but a series of perception studies showed both the initial accent (ia) and final accent (fa) to be metrically strong, independent from phrase boundaries (e.g. Astésano et al., 2012; Garnier et al., 2016; Garnier, 2018). Furthermore, in two studies directly addressing the perception of fa, participants showed little difficulty recognizing whether or not words were marked with the primary stress (Michelas et al., 2016, 2018), contradicting the notion of ‘stress deafness’ for French. Finally, the ‘optional’, secondary stress (ia) has been shown to not only be readily perceived but even expected by listeners (e.g. Jankowski et al., 1999; Aguilera et al., 2014; Astésano, 2017). That is, perception studies have shown ia to be perceived even when its phonetic correlates are suppressed or when its *f*_0_ rise peaks further along on the word (Astésano et al., 2012), pointing towards a metrical expectation for the accent. Moreover, results from a recent MisMatch Negativity study investigating the representation of ia not only provided additional evidence against the notion of stress deafness in French but also indicated a long-term memory representation and phonological preference for ia (Aguilera et al., 2014; Astésano et al., *in prep*).

Indeed, the MisMatch Negativity component (mmn) has proven a valuable tool in the study of metrical stress processing during speech comprehension. The mmn is argued to be the prototypical component for prediction mismatching input (e.g. Näätänen et al., 2007; Garrido et al., 2009; Winkler et al., 2009; Denham & Winkler, 2017). The mmn is a pre-attentive, fronto-central negative deflection peaking around 250 ms after the detection of a regularity violation (Näätänen et al., 1997) and its amplitude is held to reflect the magnitude of the deviance from what was expected (Sussman, 2007; Näätänen et al., 2007; Sussman et al., 2014; Sussman & Shafer, 2014). Such deviance can be purely acoustic (bottom-up) or it can be a deviance from a top-down derived prediction which is based on long-term memory representations (e.g. Winkler et al., 2009; Garrido et al., 2009). In the latter case, the mmn can thus index the strength of memory traces.

mmns are typically investigated in an oddball paradigm wherein a low-probability stimulus (the oddball, or deviant) occurs within a train of high-probability stimuli (Näätänen et al., 2007). The frequently occurring standard stimuli are assumed to develop predictions that are subsequently violated by the infrequently occurring deviant stimulus. The standard and deviant stimuli will usually be very similar acoustically, contrasting only on the phonological property of interest in the investigation (e.g. phoneme or stress pattern). The mmn is then obtained by subtracting the erp elicited by the standard from the erp elicited by the deviant. Therefore, the mmn represents the difference between the neural response to the frequently occurring standard stimulus and the infrequently occurring deviant stimulus, i.e. the mmn is a ‘difference wave’ that reflects the status of the manipulated phonological feature.

Importantly, whereas an mmn may be elicited by a purely acoustic difference, many studies will additionally switch the position of the deviant stimulus and the standard stimulus, such that they have another condition, wherein the deviant is presented frequently, while the (formerly) standard stimulus is presented infrequently (e.g. Honbolygó & Csépe, 2013; Aguilera et al., 2014; Scharinger et al., 2016, see also Astésano et al. *in prep*). If the standard and deviant stimuli differ only acoustically, the mmns in both conditions should be similar. Often, however, mmn amplitudes will differ, presumably due to a more established representation for one type of stimulus over the other. That is, repeatedly presenting a stimulus with a firm phonological representation, only builds on its probability leading to a large mismatch response when its anticipation is violated. In the reverse situation, when a train of improbable standards is interrupted by a more probable deviant, the violation, and thus the mismatch response, is much smaller. So, switching the positions of the standard and deviant stimuli allows for more substantial inferences on the phonological or long-term memory foundation of the manipulated phonological entity (e.g. Winkler et al., 2009; Garrido et al., 2009) and, as such, the mmn has had a substantial contribution in investigations of underspecification of phonemic representations (e.g. Eulitz & Lahiri, 2004; Näätänen et al., 2007; Winkler et al., 2009; Deguchi et al., 2010; Ylinen et al., 2016; Scharinger et al., 2016, 2017), as well as the phonological representation of stress patterns (e.g. Ylinen et al., 2009; Honbolygó et al., 2004; Honbolygó & Csépe, 2013; Aguilera et al., 2014; Honbolygó et al., 2017; Garami et al., 2017).

For instance, Honbolygó et al. (2004) investigated processing difficulties of stress patterns in Hungarian participants. The standard in their oddball study was a disyllabic word with trochaic stress, the typical stress pattern in Hungarian, while the deviant carried an iambic stress pattern. The deviant elicited two different mmns: one in response to the lack of the typical and expected stress on the first syllable, and another to the atypical additional stress on the second syllable. In a follow-up study, the trochaic and iambic stress pattern served both as standards and deviants in two separate blocks (Honbolygó & Csépe, 2013). Again, the results indicated that the deviant with an iambic stress pattern elicited two consecutive mmns, however, when the trochaic patterns had been the deviant, no mmn followed. The authors argued that the unfamiliar iambic stress pattern mismatched both the short and long-term memory representations, and, therefore, elicited the mmns, while the typical (and thus expected) trochaic stress pattern did not elicit any mmn because it did not mismatch the long-term memory representation of word stress in Hungarian. These findings provide evidence that the processing of stress pattern changes relies on language-specific long-term memory representations which may be revealed in mmn investigations (see also Ylinen et al., 2009, for similar results).

As mentioned above, in a study addressing the phonological status of the French initial accent (ia), Aguilera et al. (2014) showed that ia is not only perceived, but anticipated by listeners as belonging to the abstract representation of the word (see also Astésano et al., *in prep*). The authors manipulated the phonetic realization of ia on trisyllabic words in an oddball paradigm. Participants either listened to a version of the oddball-task wherein the stimulus +ia was in the standard position and the word −ia in the deviant position, or a version wherein ±ia positions were reversed. All listeners completed two tasks, one passive task during which they listened to the stimuli while attending a silent movie, and one active task during which the listeners were asked to respond as quickly and accurately as possible when they detected the deviant stimulus. Results indicated that the listeners clearly distinguished between the trisyllabic words carrying ia and those that did not. This again indicates that French listeners are in fact not deaf to stress, but readily perceive the accentual manipulation. Interestingly, the authors additionally observed an asymmetry between the mmn elicited by +ia deviants and the mmn following −ia deviants. That is, when the deviant had been presented without initial accent, a clear mmn component emerged, while this mmn was significantly smaller when the deviant was presented with initial accent. Not finding an mmn when presenting the oddball with ia indicates a long-term representation for the initial accent. Indeed, it is plausible that, if ia is part of a preferred stress template, only rarely presenting the template might make it the deviant within the experiment, but it does not make the template improbable. In other words, in the condition in which the oddball was presented with ia, while atypical in the context of the test, the oddball was still the expected stress template. Therefore, no mmn emerged.

In order to further ascertain that the observed mmns were independent from differences in acoustic processing, Aguilera and colleagues carried out an additional analysis wherein they compared the mmns resulting from the difference wave between −ia–deviants and −ia–standards to the difference wave between +ia–deviants and +ia–standards (i.e. between participants comparison). Again, results indicated that the difference between stimuli without initial accent was significantly larger than the difference between stimuli with initial accent, allowing for the purely acoustic interpretation of the results to be ruled out. Finally, the behavioral results from the active task confirmed the interpretation of the erp results. That is, the deviant stimuli −ia were slower to detect than the deviant stimuli +ia, and generated more detection errors. Overall, Aguilera and colleagues thus not only show that stimuli without ia are noticed by listeners, but also that ia is anticipated and attached to the metrical template underlying the representation of words.

In the current study, we set out to build on these findings and investigated the phonological representation of the French final accent in an oddball paradigm (fa). Following Di Cristo, we argue words to be encoded with bipolar stress templates underlying their representation, marking not only the left (ia) but also the right (fa) lexical boundary. Here we sought to determine whether fa is phonologically represented, similar as ia, and manipulated the presence of fa on trisyllabic words in an auditory oddball paradigm. In a first study (*Experiment 1*), participants took part in an oddball paradigm wherein either the standard word was presented with final accent and the deviant was presented without, or vice versa. In a second study (*Experiment 2*), standards were presented with their full bipolar stress templates, including both ia and fa, while deviants were either presented without fa or without ia. We expected that, if words are encoded with both accents underlying their phonological representation, then ±fa deviants should result in asymmetrical *mmn*s, similar as ±ia deviants.

### 1.1 Experiment 1: Final Accent

#### 1.1.1 Methods: Final Accent

##### Participants

The study was conducted in accordance with the Declaration of Helsinki. 21 French native speakers, aged 19 – 31 (mean age 24.0), gave their informed consent and volunteered to take part in the study. All subjects were right-handed, with normal hearing abilities and no reported history of neurological or language-related problems. Two subjects were excluded from the eeg-analysis due to excessive artifacts in the signal.

##### Speech stimuli

Two trisyllabic French nouns were used in the current experiment (‘casino’ ([kazino], *casino*) and ‘paradis’ ([paʁadi], *paradise*). The stimuli were extracted from sentences spoken by a naïve native speaker of French. Stimuli with the most natural fa (+fa) (i.e. a third syllable that was minimally 25% longer in duration than the preceding unaccented medial syllable, the primary phonetic parameter of fa Astésano, 2001) were selected by a panel of three experts. The metrical condition (±fa) was created by shortening the duration of the third syllable (fa) of the target word such that it approximated the medial, unaccented syllable and did not end in a final rise of *f*_0_ (the two main phonetic signatures of fa). This procedure was first performed automatically using a customized script in PRAAT (Boersma & Weenink, 2016) which cut the waveform, and then fine-tuned manually to correct perceptual bursts. Further, in order to keep a natural sound to the stimuli, an additionally fade-out was applied by filtering the end of the sound files with the latter half of a Hanning window.

In order to avoid mmns reflecting purely durational difference between the stimuli (i.e. total word length −fa being shorter than the total length of words +fa) (e.g. Jacobsen & Schröger, 2003; Colin et al., 2009; Honbolygó et al., 2017) and make sure the mmns had similar onset latencies between the metrical conditions, durations were equalized between ±fa stimuli by shortening the first two syllables of +fa stimuli. To additionally avoid confounds from shortening the two initial syllables, these first two syllables were shortened below the perceptual threshold following Rossi (1972) and Klatt (1976). To verify that the durational modulations on the first two syllables were not perceptible, two independent French phonetic experts completed an XO-task wherein they listened to word pairs that were either both manipulated on the first two syllables (25%), both without the durational manipulation (25%), or one with and the other without (50%). The listeners judged whether the two words were identical or different. Only stimuli with accuracy rates that were at or below chance-level were admitted in the current corpus (see figure 1 for an overview of the stimuli for both the current and second oddball study and table 1 for an overview of acoustic properties). This led to total word durations of 503 ms for ‘casino’ and 460 ms for ‘paradis’, with third syllable durations of 233.3 ms and 178.8 ms for ‘casino’ +fa and −fa, and 225.9 ms and 142.3 ms for ‘paradis’ +fa and −fa (see also table 1).

**Figure 1:**
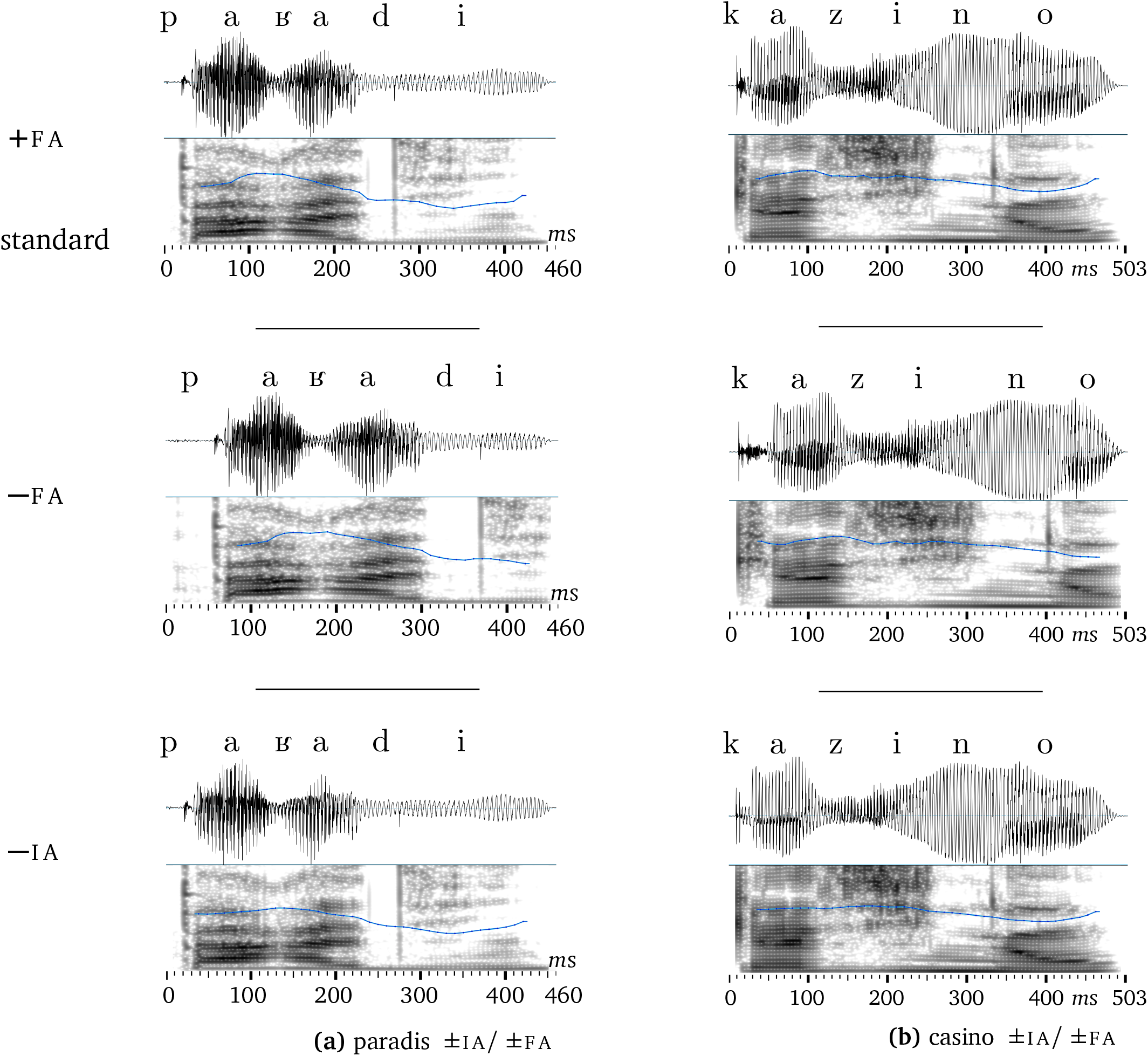
paʁadi (‘paradis’, left) and [kazino] (‘casino’, right) +fa/standard (top) and −fa (middle) and −ia (bottom). The waveforms and associated pitch tracks show how syllable duration was shortened substantially for the final syllable, and moderately for the initial two syllables.

**Table 1:** Overview of durational and *f*_0_ values, plus the timing of the third syllable (holding ±fa) onset for both ‘casino’ and ‘paradis’. Note that the +fa and −fa stimuli could serve as standards or deviants in *Experiment 1*, while, in *Experiment 2*, +fa stimuli always served as standards and −fa and −ia stimuli always served as deviants (further presented below).

|  | Total duration |  | 1st syllable |  | 2nd syllable |  | 3rd syllable |  | 3rd syllable |
| --- | --- | --- | --- | --- | --- | --- | --- | --- | --- |
|  | *ms* | *f*<sub>0</sub> | *ms* | *f*<sub>0</sub> | *ms* | *f*<sub>0</sub> | *ms* | *f*<sub>0</sub> | onset |
| CASINO |  |  |  |  |  |  |  |  |  |
| +FA/standard | 503.0 | 235.2 | 112.9 | 252.8 | 157.1 | 243.1 | 233.3 | 219.2 | 290.8 |
| −IA | 503.0 | 231.7 | 112.9 | 241.4 | 157.1 | 241.7 | 233.3 | 218.8 | 290.8 |
| −FA | 503.0 | 238.8 | 146.9 | 250.5 | 201.8 | 243.3 | 178.8 | 218.2 | 351.4 |
| PARADIS |  |  |  |  |  |  |  |  |  |
| +FA/standard | 460.0 | 212.9 | 128.7 | 245.0 | 111.6 | 233.9 | 225.9 | 185.0 | 243.1 |
| −IA | 460.0 | 208.5 | 128.7 | 228.4 | 111.6 | 225.3 | 225.9 | 186.0 | 235.0 |
| −FA | 460.0 | 224.0 | 168.8 | 244.7 | 148.3 | 239.5 | 142.3 | 189.9 | 317.5 |

Because in mmn studies which set out to investigate word processing, it is generally recommended to reduce stimulus variation between the standard and deviant as much as possible (Pulvermüller & Shtyrov, 2006; Honbolygó & Csépe, 2013), the oddball paradigm in the current study either presented only ‘casino’, or only ‘paradis’. However, because we were interested in the phonological representation of fa, which should be similar between the two words, the data obtained from both versions are combined in the analysis (see below).

In both versions, there were a total of 1092 presentations, 986 standards and 106 deviants. The deviant could be either −fa with +fa as standard, or +fa as deviant and −fa in standard position. This means that there were a total of four versions of the oddball paradigm: (1) casino–deviant +fa, (2) casino–deviant −fa, (3) paradis–deviant +fa, and (4) paradis–deviant −fa.

##### Procedure

Each participant was comfortably seated in an electrically shielded and sound attenuated room. Stimuli were presented through headphones using Python2.7 with the PyAudio library on a Windows XP 32-bit platform. To ensure attention was diverted from the stimuli, participants watched a silent movie with no text (*Best of mr. Bean*).

Lists were assigned randomly: 4 participants listened to the casino-deviant +fa version, 3 listeners to the casino-deviant −fa version, 7 participants listened to paradis-deviant +fa and finally 5 participants had the paradis-deviant −fa version. This meant that data was obtained from 11 participants for the version in which +fa stimuli were in deviant position and −fa stimuli in standards, and from 8 participants for the version wherein ±fa positions were reversed.

Each participant listened to the complete list of 1092 stimuli (986 standards, 106 deviants) in one block, which lasted approximately 25 minutes. Deviants were interspersed randomly and online, while avoiding two consecutive occurrences and making sure that each list started with at least 25 standards. Finally, the inter-trial interval (iti) consisted of stimulus duration plus an inter-stimulus interval (isi) of 600 ms.

##### EEG recording and preprocessing

eeg data were recorded with 64 Ag/AgCl-sintered electrodes mounted on an elastic cap and located at standard left and right hemisphere positions over frontal, central, parietal, occipital and temporal areas (International 10/20 System; Jasper, 1958). The eeg signal was amplified by BioSemi amplifiers (ActiveTwo System) and digitized at 2048 Hz.

The data were preprocessed using the `EEGlab` package (Delorme & Makeig, 2004) with the `ERPlab` toolbox (Luck et al., 2010) in Matlab (Mathworks, 2014). Each electrode was re-referenced offline to the algebraic average of the left and right mastoids. The data were band-pass filtered between 0.01–30 Hz and resampled at 128 Hz. Following a visual inspection, signal containing EMG or other artifacts not related to eye-movements or blinks was manually removed. Independent Components Analysis (ica) was performed on the remaining data in order to identify and subtract components containing oculomotor artifacts. Finally, data were epoched from −0.2 to 1 seconds surrounding the onset of the stimulus and averaged within and across participants to obtain the grand-averages for each of the two stress conditions.

##### Analysis

The method of eeg provides high temporal precision. However, the high temporal resolution comes at the cost of many comparisons when erp amplitude values for each individual electrode, at each recorded time-point, are tested independently, using standard parametric statistics (e.g. ANOVA). Because EEG measures are not independent, but instead temporally and spatially correlated, we use a non-parametric *t*_max_ permutation test to analyze the data (Groppe et al., 2011; Luck, 2014) using the `Mass Univariate ERP Toolbox` (Groppe et al., 2011) in Matlab (Mathworks, 2014). We were interested in modulations of the mmn as elicited by the presence/absence of fa and therefore specifically tested for differences in the time-window between 551 −651 ms. Furthermore, because the mmn is a fronto-centrally located deflection we selected the fronto-central electrodes (Fz, Cz, FC1, FC2, F3, F4, C3 and C4) for the statistical analyses. Each comparison of interest was analyzed with a separate repeated measures, two-tailed *t*-tests, using the original data and 2500 random permutations to approximate the null distribution for the customary family-wise alpha (*α*) level of 0.05.^2^

#### 1.1.2 Results

Similar to Aguilera et al. (2014), we obtained no significant mmn when the deviant had been +fa (critical *t*-score: ±4.3095, *p* = 0.8396, *ns*). This indicates that even though the +fa stress template was rare in the experimental setting, listeners *still* expected words to be marked with final accent. Presenting the deviant without final accent elicited a marginally significant mmn (critical *t*-score: ±4.2958, *p* = 0.0652). Visual inspection suggests the mmn was located at left frontal electrodes, starting 600 ms post stimulus onset (i.e. ~ 300 ms post deviance detection). Furthermore, we observed an asymmetry between mmns; the mmn was significantly more ample when the deviant had been presented −fa than when it had been presented +fa (critical *t*-score: ±3.1505, *p* < 0.05, see figure 2).

**Figure 2:**
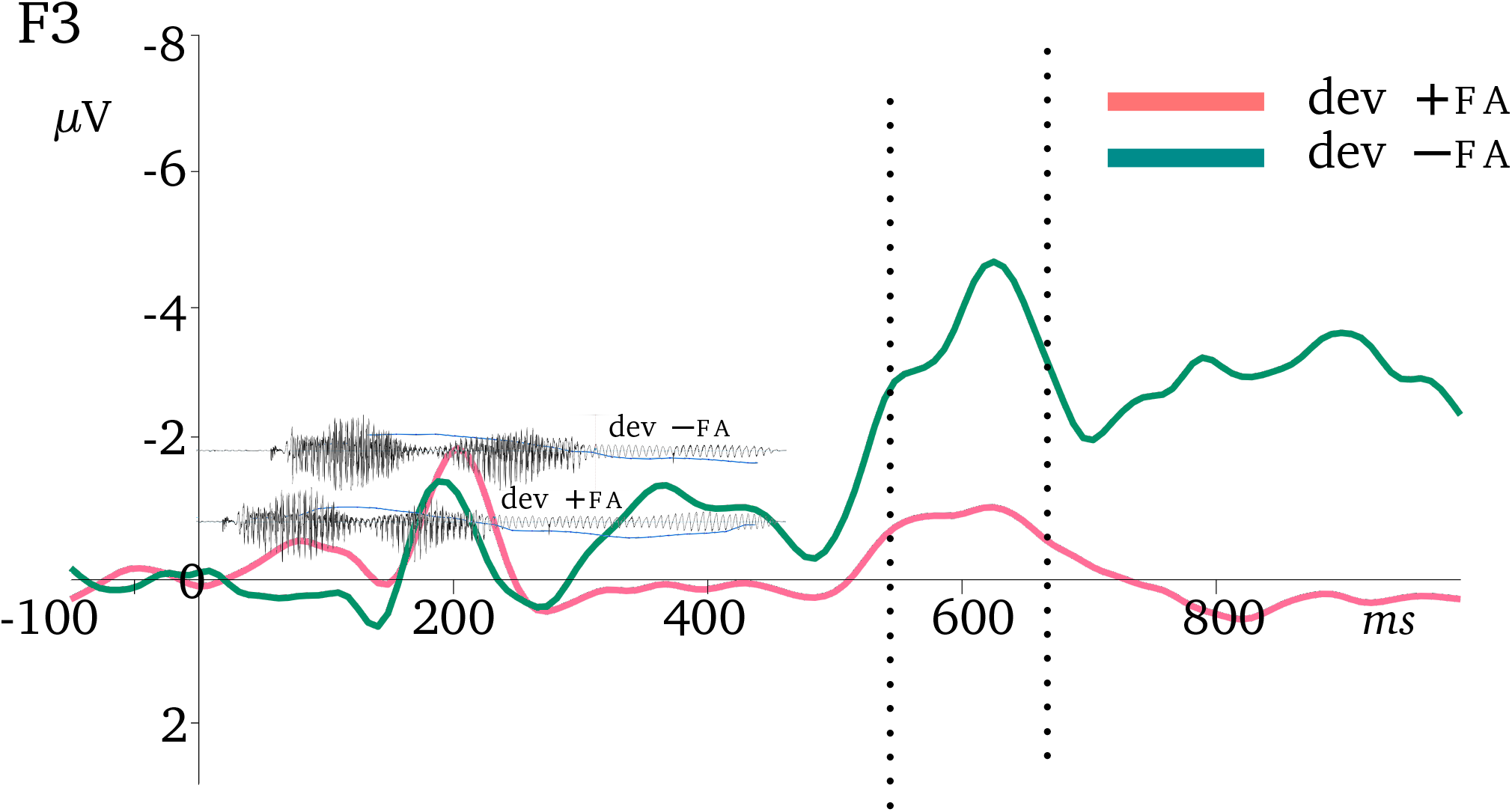
mmn components for +fa (in pink) and −fa (in green) deviants, recorded at the F3 (left frontal) electrode, with the oscillogram of the deviant stimuli [paKadi] plotted in the background. Waveforms and oscillograms are temporally aligned to indicate the relation between the offset of the ±fa manipulation and the resulting stimulus-locked event-related potentials. The tested time-window is indicated by dashed vertical lines. For ease of presentation, erp waveforms are low-pass filtered at 10 Hz and negativity is plotted as an upward deflection.

Finally, in the comparison between participants (i.e. comparing identical stimuli that differed in position within the oddball experiment) there was a significant difference between +fa in deviant position versus +fa in standard position at frontal (F4) and central (C4) electrodes during the whole time-window (critical *t*-score: ±3.7416, *p* < 0.05), while there was no such difference for stress templates −fa (critical *t*-score: ±4.394, *p* = 0.84, *ns*, see figure 3).

**Figure 3:**
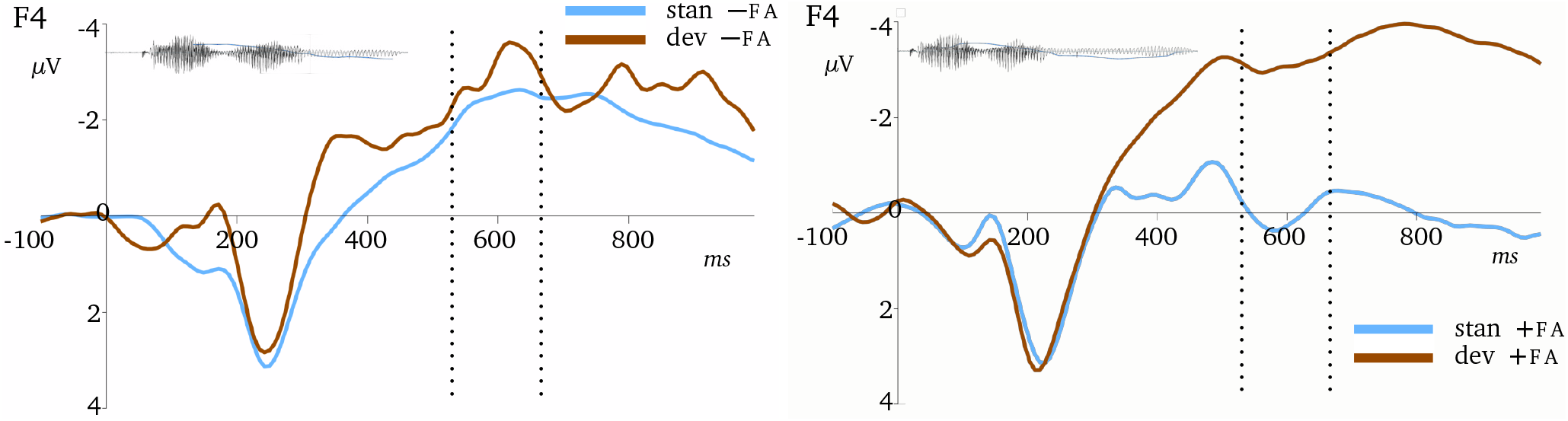
erp components for −fa (left) and +fa (right) stimuli, recorded at the F4 (right frontal) electrode, with the oscillograms of [paKadi] plotted in the background to indicate the relation between the offset of the ±fa manipulation and the resulting stimulus-locked erps. The tested time-window is indicated by dashed vertical lines. For ease of presentation, erp waveforms are low-pass filtered at 10 Hz and negativity is plotted as an upward deflection.

Note that the results presented here partially contradict those reported in Aguilera et al. (2014) in which ia had been manipulated. In Aguilera et al. (2014), the between listeners analysis demonstrated a bigger difference between standards and deviants when stimuli had been presented −ia, than when they had been presented +ia. This discrepancy potentially indicates differential processing between ia and fa, which is elaborated upon in the main discussion of the two experiments.

### 1.2 Experiment 2: Initial and Final Accent

#### 1.2.1 Methods: Initial and Final Accent

##### Participants

20 French native speakers, aged 19 – 45 (mean age 23.7), took part in the study. None of the participants had taken part of the previous mmn study and all were right-handed, with normal hearing abilities and no reported history of neurological or language-related problems. Each of the participants gave their written consent and was paid a small fee for their participation.

##### Speech stimuli

The same two trisyllabic French nouns used in the previous study, were used in the current experiment (‘casino’ ([kazino], *casino*) and ‘paradis’ ([paʁadi], *paradise*). The natural ia (+ia) was resynthesized without ia (−ia) using a customized quadratic algorithm in Praat (Boersma & Weenink, 2016). Using the same algorithm as in Aguilera et al. (2014), the *f*_0_ value of the first vowel (i.e. ia) was lowered near the *f*_0_ value of the preceding (unaccented) determinant, to de-accentuate the first syllable (i.e. remove ia). The algorithm progressively modified the *f*_0_ values to reach the *f*_0_ value at the beginning of the last (accented) vowel. This quadratic transformation allowed for micro-prosodic variations to be maintained, thus keeping the natural sound of the stimuli. The +ia stimuli were forward and back transformed to equalize the speech quality between +ia and −ia stimuli (see Aguilera et al., 2014, for more information on the manipulation of ia). Refer to figure 1 and table 1 for an overview of stimuli properties for both words ±ia and ±fa. As in *Experiment 1*, we presented lists either only with ‘casino’, or only with ‘paradis’. However, because we were interested in the phonological representation of the accent (whether ia or fa), which should be similar between both words, the data obtained from both versions are again merged in the analysis.

##### Procedure

Each participant was comfortably seated in an electrically shielded and sound attenuated room. Stimuli were presented through headphones using Python2.7 with the PyAudio library on a MacOS Sierra platform. Similar as in the previous experiment, participants watched a silent movie to ensure their attention was diverted from the stimuli. Each participant listened to all 1200 stimuli (1000 standards, 100 deviants −ia, 100 deviants −fa) in one block, which lasted for approximately 25 minutes. Deviants were interspersed randomly and online, while avoiding two consecutive occurrences of the same deviant and making sure that each list started with 25 standards. Finally, the same inter-trial interval (iti) was used as in the previous oddball study, and consisted of stimulus duration plus inter-stimulus interval (isi) of 600 ms.

##### EEG recording and preprocessing

eeg data were recorded with 64 Ag/AgCl-sintered electrodes mounted on an elastic cap and located at standard left and right hemisphere positions over frontal, central, parietal, occipital and temporal areas (International 10/20 System; Jasper, 1958). The eeg signal was amplified by BioSemi amplifiers (ActiveTwo System) and digitized at 2048 Hz. The data were preprocessed using the `EEGlab` package (Delorme & Makeig, 2004) with the `ERPlab` toolbox (Luck et al., 2010) in Matlab (Mathworks, 2014). Each electrode was re-referenced offline to a common average reference. The data were band-pass filtered between 0.01 −30 Hz and resampled at 256 Hz. Following a visual inspection, signal containing EMG or other artifacts not related to eye-movements or blinks was manually removed. Independent Components Analysis (ica) was performed on the remaining data in order to identify and subtract components containing oculomotor artifacts. Finally, data were epoched from −0.2 to 1 seconds surrounding the onset of the stimulus and averaged within and across participants to obtain the grand-averages for each of the stress conditions.

##### EEG analysis

The data were analyzed with the non-parametric *t*_max_ permutation test (Groppe et al., 2011; Luck, 2014) using the `Mass Univariate ERP Toolbox` (Groppe et al., 2011) in Matlab (Mathworks, 2014). We were interested in modulations of the mmn as elicited by the presence/absence of ia and fa. Therefore, we specifically tested for differences in the time-windows between 201–301 ms and 551–651 ms, respectively. Furthermore, because the mmn is a fronto-centrally located deflection we specifically tested the Fz, Cz, FC1, FC2, C3, C4, F1, F2, F5h, F6h and FCz electrodes in both time-windows. Each comparison of interest was analyzed with a separate repeated measures, two-tailed *t*-tests, using the original data and 2500 random permutations to approximate the null distribution for the customary family-wise alpha (*α*) level of 0.05.

#### 1.2.2 Results

Both −ia and −fa deviants elicited a mmn, although the mmn was smaller, and only marginally significant, when the deviant had been −ia (critical *t*-score: ±3.368, *p* = 0.06) compared to when the deviant had been −fa (critical *t*-score: ±3.4322, *p* < 0.05) (see figure 4). This difference is interpreted in the main discussion of both oddball studies below.

**Figure 4:**
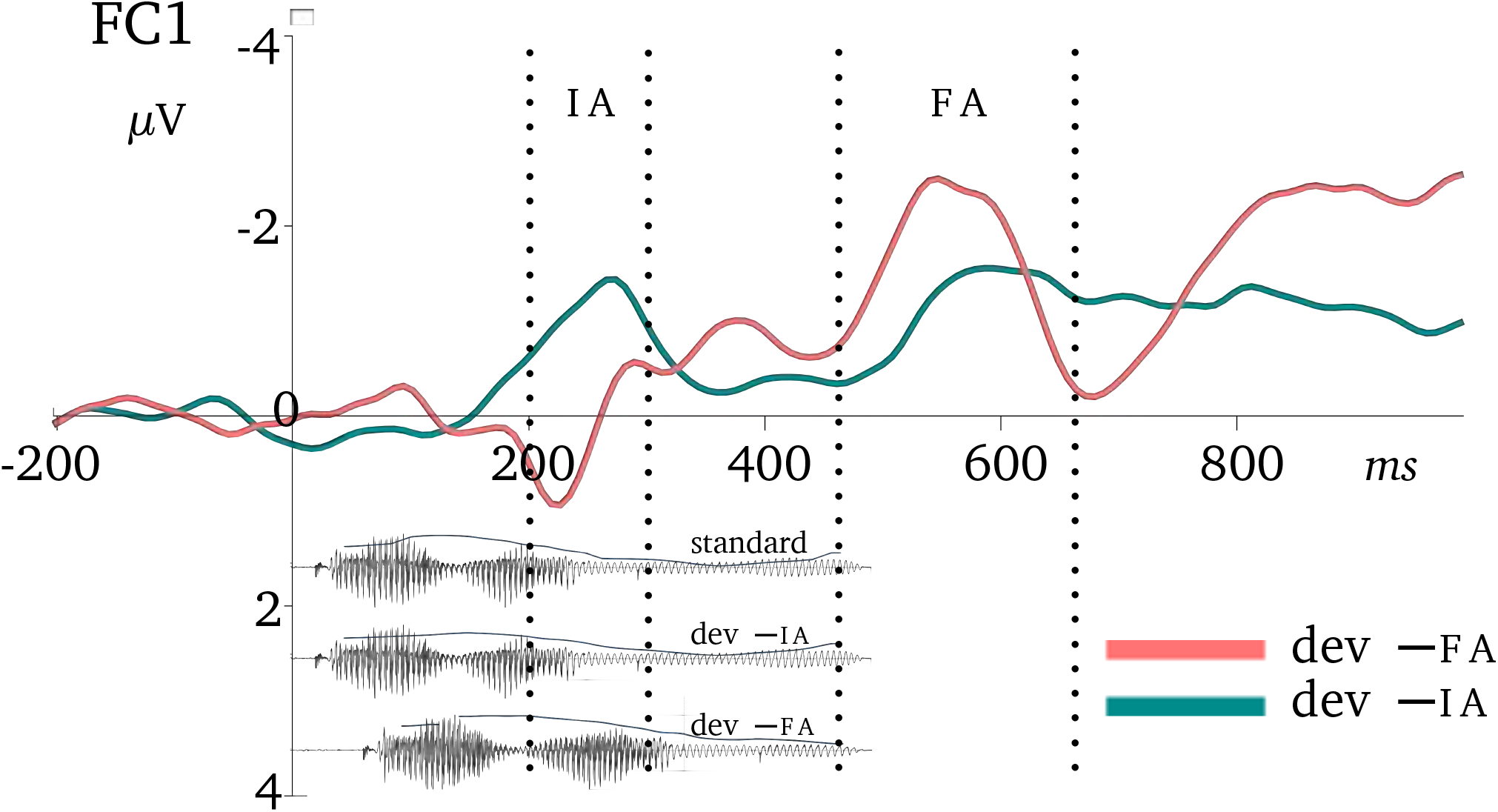
mmn components for −ia (in green) and −fa (in pink) deviants, recorded at the FC1 (left frontal) electrode, with the oscillogram of all stimuli type of [paʁadi] plotted in the background. Waveforms and oscillograms are temporally aligned to indicate the relation between the offset of the ±ia and ±fa manipulation and the resulting stimulus-locked event-related potentials. Tested time-windows are indicated by dashed vertical lines. For ease of presentation, erp waveforms are low-pass filtered at 10 Hz and negativity is plotted as an upward deflection.

## 2 Discussion

In the current studies, we sought to investigate the phonological representation of French accentuation. We took advantage of the mmn component, which is held to index the strength of memory traces underlying phonological information. We based our expectations on the results presented in Aguilera et al. (2014), which had previously shown the French secondary initial accent (ia) to be encoded in long-term memory and to be expected by listeners.

We were first specifically interested in the representation of the French primary final accent (fa) and manipulated its presence on trisyllabic words in an auditory oddball paradigm. There were two versions of this paradigm; in one version the standards were presented with fa, while the deviants were presented without fa, and in the other version, positions were reversed. As we will discuss in more detail below, our results showed a pre-attentive expectation for words to be presented with final accent and a general dispreference for words presented without the accent. However, our results partially deviated from those obtained in Aguilera et al. (2014), i.e. while the asymmetrical mmns elicited by ±fa deviants are congruent to the results reported in Aguilera et al. (2014), the comparisons between participants differed. In order to better understand this deviance between ia and fa, in *Experiment 2*, we orthogonally manipulated the presence of both the final accent and the initial accent within the same paradigm. That is, in this second experiment, both −fa and −ia stimuli served as deviants, while the standard was consistently presented with both the final and initial accent. We obtained mmn difference waves to both −ia and −fa deviants. The amplitudes of the respective mmns, however, differed in size, possibly reflecting a different functional role for the accents in their marking of the word.

Below, we will discuss our findings in turn: In section 2.1, we present the results obtained in the first oddball study that show fa to not only be readily perceived, but also to be expected by the listener and phonologically natural. In section 2.2, we discuss the differential processing of −fa and −ia deviants. We will interpret the results from an acoustic, exogenous point of view, as well as inspect the possibility for this difference to reflect more substantial, endogenous differences in the functions of the respective accents during word processing.

### 2.1 Phonological representation of the French final accent

In *Experiment 1*, wherein we had concentrated on the representation of the French final accent, we observed an asymmetry between mmns elicited by deviants presented without final accent, compared to those elicited by deviants that had been presented with final accent. More specifically, the mmn was significantly more ample when the deviant had been presented −fa, than when it had been presented +fa (see figure 2). This asymmetry indicates that the final accent is encoded in long-term memory and part of the expected stress template.

Our comparisons between participants corroborate with this interpretation (see figure 3). Presenting words without final accent elicited an ample erp deflection, *irrespective* of the position of the stimuli within the experimental setting. That is, words without final accent appeared to require more cognitive effort, *regardless* of whether they had been the standards or the deviants in the oddball paradigm. This result shows stress templates without fa to be generally disfavored. Indeed, if there had been no preference for one stress pattern over the other, then repeatedly presenting words without final accent (i.e. when −fa is in standard position) *should* have made the stress pattern −fa the probable stress template. Clearly, it did not; even in standard position, the stress pattern without final accent remained unexpected. In other words, listeners continued to anticipate words to be marked with final accent, most likely due to its established phonological representation.

The comparison between standards and deviants presented with final accent points to the same conclusion. In this comparison, position within the experimental setting *did* matter. Recall that the mmn may reflect both a prediction error when anticipations based on established phonological representations are violated, as well as a mismatch within the experimental context. In the comparison between words +fa, interrupting a train of −fa stimuli with the sudden presentation of a template with fa, elicited a small prediction error, while no such prediction error followed the final accent when +fa stimuli were in the standard position. This finding again disproves the notion of stress deafness, i.e. listeners *readily notice* the accent when deviants +fa contrasted to the train of −fa templates. In other words, listeners detected fa when it mismatched the short-term anticipation established by the repeated stress templates −fa, negating their alleged phonological deafness. However, as was explained above, the mismatch did *not* result in a significant mmn (far from it, see figure 2), because presenting stimuli without final accent, even when congruent within the experimental setting (i.e. when −fa was in the standard position), remained unexpected due to the long-term phonological representation of the final accent.

In sum, we show that the final accent is readily perceived and elicits a small prediction error when it mismatches short-term memory, while stress patterns without final accent mismatch both short-*and* long-term memory representations and thus do not appear to be the expected metrical pattern in French.

### 2.2 Differential processing between the initial and final accents

While the asymmetrical mmns elicited by ±fa deviants are congruent to the results reported in Aguilera et al. (2014), the comparisons between participants differed. Where Aguilera and colleagues obtained a bigger difference wave after words were presented without ia underlying their stress template, even when comparing acoustically identical stimuli in both standard and deviant position, we obtained results opposite to that (i.e. there was a bigger difference between +fa stimuli than between −fa stimuli). To better understand this incongruence, in *Experiment 2*, we orthogonally manipulated the presence of both the final accent and the initial accent within the same paradigm. That is, in this second study, both −fa and −ia stimuli served as deviants, while the standard was consistently presented with both the final and the initial accent.

We obtained two consecutive mmn deflections, one reflecting the absence of ia, the other reflecting the absence of fa (see figure 4). The amplitudes of the mmns were, however, different in size, with the mmn following deviants −fa being more ample than the mmn following deviants −ia. These results could inform us about differences in the strength of the memory representations between ia and fa, with the final accent holding a stronger memory trace and being anticipated to a greater extent by listeners than the initial accent. However, there are several alternative explanations which are also compatible, and, possibly, more likely explain the different mmns: one reflecting a purely exogenous, acoustic interpretation, and the other involving a more substantial, endogenous difference in the accents’ respective functions during speech processing. Both accounts are discussed below.

#### 2.2.1 Exogenous interpretation

In the exogenous interpretation, the dissimilar mmn amplitudes between ia stimuli and fa stimuli reflect differences in acoustic processing. Indeed, the acoustic manipulations had not been the same between our ±ia and ±fa stimuli, the former involving exclusively a manipulation of the *f*_0_ rise, and the latter involving mainly a durational change. It is possible that French listeners are more sensitive to durational changes than to changes in pitch movement (see e.g. Partanen et al., 2011, for an mmn study showing just that for Finnish speakers, although also note that sensitivity to stress phonetic features is likely language specific). Moreover, while the presence of ia was *only* manipulated in *f*_0_, the durational change of fa led to the additional disappearance of the accent’s final rise (see figure 1), the secondary phonetic characteristic of fa. This means that stimuli without fa differed from stimuli with *fa* on two acoustic parameters, while ±ia stimuli differed only in *f*_0_. Because mmn amplitudes are held to reflect the *magnitude* of the deviance between standard stimuli and deviants (Sussman, 2007; Näätänen et al., 2007; Sussman et al., 2014; Sussman & Shafer, 2014), these exogenous interpretations may at least in part explain the observed mmn differences between our −ia and −fa stimuli.

However, a purely acoustic interpretation less straightforwardly accounts for the different findings in the between participants comparisons observed in the current study versus those presented in Aguilera et al. (2014). Therefore, we consider it more likely that the dissimilar amplitudes reflect different respective roles for the accents during speech processing. Indeed, while the initial accent sits at the left word boundary and, as such, could signal word onsets and cue listeners on when to initiate lexical access, the final accent, which is located at the right word boundary, likely holds different functions, such as marking the word’s offset and cue listeners on when to finalize their analysis of the word. In this view, the respective mmns then reflect different interactions between the accents and the stages in speech perception, which we will turn to next.

#### 2.2.2 Endogenous interpretation

Speech perception unfolds in three stages: an acoustic stage, during which the speech signal is spectrally decomposed and distinguished from non-speech sounds, a pre-lexical stage, during which phonological information is assembled and matching lexical candidates are activated, and, finally, the lexical stage, wherein candidates compete and are evaluated up until one word can be selected for word recognition. In our view, the initial accent is more likely to interplay with the pre-lexical stage during which lexical hypotheses are derived and activated, while the final accent will presumable be more involved in the later lexical stage which ends in the recognition of the word. In terms of the Cohort model (Marslen-Wilson & Welsh, 1978; Wilson, 1990), the initial accent (the word’s earliest phonological information) activates similar lexical representations into the, so-called, cohort. As the speech signal continues, matching candidates are additionally activated while, when words without final accents seize to match the activated representations, these are disregarded from the cohort or lessened in activation levels. In other words, the initial accent plausibly has more effect on the *start* of the process of word recognition and on early lexical activation levels, while the final accent is more likely involved in the *outcome* or *wrap-up* of the lexical competition.

Note that, in this view, dissimilar mmns elicited by ±ia versus ±fa stimuli are not only explained in terms of different interactions during the process of word recognition, but also in terms of the precision of the prediction to which the stress patterns are compared. According to the theory of predictive coding, predictions which are precise require less additional cognitive effort than predictions which are more generic. Stimuli without ia differ from the prediction phonologically, i.e. the listener has a general phonological preference or expectation for words to be presented with both ia and fa in their underlying stress templates. When the deviance is however later in the word, as with −fa, the listener’s prediction more pointedly concerns the phonological stress template marking the right boundary of the particular lexical item expected from the train of standards. That is, one can imagine that, if fa cues the lexical offset, listeners could have imagined words without final accent to be part of, or embedded in, a longer word, therefore deleting the anticipated word boundary. Indeed, words can have other words partially or wholly embedded within them, such that the speech stream usually matches with multiple lexical candidates (*the embedding problem*, e.g. ‘paradis’ is a word on its own, but can also be at the onset of, for example, ‘paradisiaque’ or ‘paradigmatique’). When presented stress patterns mismatch the expected stress template, this can lead to wrongfully deleting a word boundary. Indeed juncture misperception studies on English and Dutch, languages wherein stress is often word-initially (Cutler & Carter, 1987; Vroomen & de Gelder, 1995), have shown listeners to erroneously insert a word boundary when encountering a strong syllable (for instance, “analogy” → “an allergy”) or delete a word boundary before a weak syllable (for instance, “my gorge is” → “my gorgeous”) (e.g. Cutler & Butterfield, 1992; Vroomen et al., 1996).

In fact, also French listeners have been found to segment speech on fa in ambiguous sentences (see e.g. Banel & Bacri, 1994; Bagou et al., 2002; Christophe et al., 2004, for studies wherein fa signaled the right phrase boundary). For example, Banel & Bacri (1994) found listeners to use the lengthened syllables as a right boundary cue and, consequently, segmented immediately after them. That is, when listeners were asked to interpret ambiguous speech sounds such as [bagaʒ] which may be segmented into two words ‘bas + gage’ (low + pledge) or can be interpreted as ‘baggage’ (luggage), listeners favored the former interpretation when the syllables were marked with a trochaic stress template (long—short), while conversely favoring the latter interpretation when the stress template had been iambic (short—long). That is, lengthened syllables encouraged a boundary to the right, while short syllables did not. Because, in French, prosodic descriptions do not include the lexical word, the boundary was attributed to the phrasal domain. However, it is possible that fa might have also cued the right lexical boundary in that study.

Similarly, in the study on the interaction between metrical structure and semantic processing, Magne et al. (2007) artificially lengthened the medial syllable. This metrical ‘incongruity’ was found to obstruct semantic processing, possibly because listeners segmented speech on the medial syllable and, thus, before the word’s actual offset. In the current study, shortening the final syllable in the deviant position, may have led the deviant to not only mismatch with the anticipated phonological stress template, but change the predicted lexical item because it was missing its right boundary mark (e.g. “paradis” → “paradigmatique”). That is, listeners may have noticed the acoustic mismatch (i.e. syllable length and *f*_0_ movement), the phonological incongruence (i.e. ±fa), and the lexical difference (‘paradis’ → ‘paradigmatique’). In other words, repeatedly presenting the same lexical item in the standard position, led to more specific anticipations and activations of lexical candidates, which, in turn, resulted in mmns reflecting the concurrent detection of several deviances: (1) the acoustic deviance, (2) the phonological mismatch and, possibly, (3) the mismatch to the lexical prediction (see e.g. Pulvermüller & Shtyrov, 2006; Jacobsen et al., 2004; Honbolygó et al., 2004; Honbolygó & Csépe, 2013; Honbolygó et al., 2017; Ylinen et al., 2009; Garami et al., 2017; Zora et al., 2016, for oddball studies investigating obstructed processing due to mismatching stress templates on words and/or pseudowords in Hungarian, Finnish and Swedish).

However, if the differences between ia and fa reflect interactions with different stages during word recognition, then, while interesting, the oddball paradigm (and mmn) unfortunately is not well suited to observe them. Clearly, oddball paradigms provide a rather artificial listening situation, wherein it is not clear whether each word presentation (whether in standard position or as deviant) encourages a fresh attempt to lexical access. That is, arguably the repeated presentation of the same word may involve a process different from normal listening situations wherein listeners go through all three stages of word recognition. Future studies adopting different paradigms that encourage lexical access (e.g. a lexical decision paradigm) may be better suited to observe the possibly differential contributions of ia and fa to the process of word recognition.

## 3 Conclusion

In sum, in this oddball study, we investigated the cognitive representation of the French accentuation. The French initial accent had previously been shown to not only be readily perceived but expected by French listeners as part of the stress pattern underlying the presented words, indicative of a functional role for the accent on the analysis of speech. The results of the present study how that fa—just as ia—is not only perceived, but anticipated by listeners. Unlike the results reported in Aguilera et al. (2014), when the standard was presented without fa, it remained unexpected, despite its high probability within the experimental context. This result suggests that the deviant without fa remained improbable within the experimental setting, indicating a longterm representation of the accent and underlining listeners’ expectation for words to be marked by stress templates which also include fa. Moreover, we observed an asymmetry between deviants presented with fa and deviants presented without, with larger mmn amplitudes when the deviant had been presented without fa. In this respect, the results are congruent to the asymmetrical mmns reported in Aguilera et al. (2014) in which ia had been manipulated, and, together, the results are in line with Di Cristo’s model, and imply a cognitive, phonological expectation for metrically strong syllables at both left and right lexical boundaries. Altogether, the results contradict the traditionally accepted view of French as a language without accent and, instead, suggest accentuation to have a functional role in word level processing.

## Acknowledgments

This study was supported by the Agence Nationale de la Recherche grant ANR-12-BSH2-0001 (PI: Corine Astésano).

## Footnotes

1 Note that while the term ‘stress deafness’, when taken literally, implies a phonological deafness for French listeners, and is in fact often interpreted as such, Dupoux et al. (1997) intended for a more nuanced interpretation, wherein speakers of languages with fixed, non-distinctive stress do not encode stress templates into their mental lexicon and are consequently less sensitive to variable, lexically distinctive stress in foreign languages.

2 In fact we used more than twice the number of permutations suggested for an alpha at 5% (Manly, 2006) so as to be even more certain of obtaining reliable results.

